# Genomic and structural features of the Yellow Fever virus from the 2016-2017 Brazilian outbreak

**DOI:** 10.1101/179481

**Authors:** Mariela Martínez Gómez, Filipe Vieira Santos de Abreu, Alexandre Araujo Cunha dos Santos, Iasmim Silva de Mello, Marta Pereira Santos, Ieda Pereira Ribeiro, Anielly Ferreira-de-Brito, Rafaella Moraes de Miranda, Marcia Gonçalves de Castro, Mario Sergio Ribeiro, Roberto da Costa Laterrière, Shirlei Ferreira Aguiar, Guilherme Louzada Silva Meira, Deborah Antunes, Pedro Henrique Monteiro Torres, Ana Carolina Paulo Vicente, Ana Carolina Ramos Guimarães, Ernesto Raul Caffarena, Gonzalo Bello, Ricardo Lourenço-de-Oliveira, Myrna Cristina Bonaldo

**Affiliations:** Laboratório de Biologia Molecular de Flavivírus, Instituto Oswaldo Cruz, Fundação Oswaldo Cruz, Rio de Janeiro, Brazil; Laboratório de Mosquitos Transmissores de Hematozoários, Instituto Oswaldo Cruz, Fundação Oswaldo Cruz, Rio de Janeiro, Brazil; Instituto Federal do Norte de Minas Gerais, Salinas, MG, Brazil; Superintendência de Vigilância Epidemiológica e Ambiental, Secretaria Estadual de Saúde, Rio de Janeiro, Brazil; Núcleo Especial de Vigilância Ambiental, Secretaria Estadual de Saúde do Espírito Santo, Vitória, Brazil; Laboratório Central de Saúde Pública Noel Nutels (LACEN-RJ), Rio de Janeiro, Brazil; Programa de Computação Científica (PROCC), Fundação Oswaldo Cruz, Rio de Janeiro, Brazil; Laboratório de Genética Molecular de Microorganismos, Instituto Oswaldo Cruz, Fundação Oswaldo Cruz, Rio de Janeiro, Brazil; Laboratório de Genômica Funcional e Bioinformática, Instituto Oswaldo Cruz, Fundação Oswaldo Cruz, Rio de Janeiro, Brazil; Laboratório de AIDS e Imunologia Molecular,Instituto Oswaldo Cruz, Fundação Oswaldo Cruz, Rio de Janeiro, Brazil

**Keywords:** Yellow fever virus, 2016-2017 Brazilian outbreak, amino acid changes, genetic diversity, evolution

## Abstract

Brazil has been suffering a severe sylvatic epidemic of yellow fever virus (YFV) since late 2016. Analysis of full-length YFV genomes from all hosts involved in the Brazilian 2017 outbreak reveals that they belong to sub-lineage 1E within modern-lineage, but display several unique amino acid substitutions in highly conserved positions at NS3 and NS5 viral proteins. Evolutionary analyses indicate that YFV carrying that set of amino acid substitution circulates in the Southern Brazilian region for several months before being detected in December 2016. Structural and selection analyses support that some of these substitutions were under positive selection and could impact enzyme structure and function. Altogether, this evidence demonstrated that the current Brazilian YFV carries unique amino acid signatures in the non-structural proteins and support the hypothesis that those substitutions may be affecting the viral fitness and transmissibility.

## INTRODUCTION

Yellow fever (YF) is a viral disease transmitted by the bite of infected mosquitoes in Africa and South America, affecting around 200,000 people annually, mostly in Africa (1-3). There are two main epidemiological cycles: the enzootic sylvatic cycle where the virus is transmitted between non-human primates (NHP) and wild arboreal mosquitoes of genus *Aedes, Haemagogus* and *Sabethes*, and in which humans can be accidentally infected, and the urban cycle where inter-human transmission is ensured by the domestic and anthropophilic mosquito *Aedes aegypti* (3). While only the sylvatic cycle has been reported in the Americas during nearly the last 75 years, in Africa people may acquire the infection in both cycles, besides in an intermediate cycle occurring in rural areas close to forests (2, 4).

The causative agent is the yellow fever virus (YFV) (genus: *Flavivirus*, family: *Flaviviridae*), presenting a single-positive-sense RNA genome, containing a 5’ end cap structure, that is translated in a single immature polyprotein precursor. The precursor polyprotein is cleaved into three structural proteins, capsid (C), envelope (E), and membrane protein (M) and seven non-structural proteins, NS1, NS2A, NS2B, NS3, NS4A NS4B, and NS5 (2). The virus was originated in Africa, where five genotypes have been documented, being two from West Africa (West Africa I and II) and three in East and Central Africa (East Africa, East/Central Africa, and Angola). The YFV virus has spread from Africa to the Americas together with the invasive mosquito *Ae. aegypti* where it evolved in the last four centuries into two genotypes (South America I and II) derived from the Western African ancestor (5-7). The South American genotype I is the most spread and frequently detected during the epizootics and epidemics waves in Brazil and other countries of South America (8, 9). Until the 1990’s, the transmission area in Brazil was primarily limited to the Amazon forest, in the Northern, and the savanna-like *cerrado,* in Center-West region. However, in about two decades, the YFV territory has progressively expanded Southward and Eastward reaching the Atlantic forest and other biomes from the country´s most populated regions(10). During this boundary expansion, five viral sub-lineages (1A to 1E) successively arose within the genotype I. They were distinguished by analysis of partial nucleotide sequencing of the YFV genome particularly the pre-membrane and envelope (prM/E) gene junction (8, 9, 11). Most recently, South America genotype I was divided into two major lineages named as Old lineages (enclosing Old Para, and 1A, 1B, and 1C sub-lineages) and Modern lineage (enclosing Trinidad and Tobago, and 1D and 1E sub-lineages) (11, 12).

A rapid expansion of the YFV area has reported since late 2016 in Southeast Brazil (Fig.1) (13, 14). From December 2016 to May 2017, the YFV quickly spread from the transition zone between the *cerrado* and the Atlantic forest inland of Minas Gerais State (MG) to the coastal areas in the Espírito Santo (ES) and Rio de Janeiro (RJ) states, then approaching to densely populated sites with insignificant vaccination coverage under influence of the rain forest. A YFV outbreak of unprecedented sanitary severity and causing an ecological disaster was recorded. In a few months, around 3,850 NHPs died, and nearly 800 human cases with 435 deaths were registered, of which 274 were YF confirmed until July 2017 (13, 14). Interestingly, during this ongoing outbreak analysis of complete nucleotide genome sequences of the YFV obtained from the blood of two howler monkeys from a single locality in ES confirmed that they cluster within the sub-lineage 1E. Furthermore, these strains revealed new polymorphisms comprising several amino acid substitutions mainly located in the components of the viral replicase complex, in the protease domain of the NS3 protein, and in the methyltransferase (MTase) and RNA-dependent RNA polymerase (RdRp) domain of the NS5 protein(15). The NS3 protein, a viral multi-functional protein, carries a chymotrypsin-like serine protease activity (NS3pro) at its N-terminal and an helicase activity (NS3hel) at its C-terminal domain(16). The NS3pro associated with its cofactor NS2B cleaves all cytoplasmic protein junction sites of the precursor polyprotein. The NS5 protein contains two functional domains with a capping related MTase and the central replication enzyme RdRp (17-19). Connecting MTase and RdRp, there is a linker of 5–6 residues (residues 266–271; NS5 Dengue virus position), which has an essential role in NS5 conformation and protein activity (20). Also, NS3 and NS5 have been associated with innate immune system evasion (21, 22). It was unclear, however, whether the observed amino acid substitutions are genetic signatures of the most recent YF outbreaks as few complete genomes of current circulating viral strains were available (8, 15).

In this study, we elucidated the complete genome sequence of 12 YFV strains from three hosts (NHPs, mosquitoes, and humans) involved in the transmission cycle of the current Brazilian outbreak in two Southeastern states (RJ and ES). Sequences were analyzed to establish whether specific amino acid changes previously observed are fixed in other recent YFV samples and therefore constitute a molecular signature of the 2017 YFV. We also created homology models for NS3 and NS5 to determine the location and potential effects of amino acid substitutions on NS3pro and NS5 proteins. Moreover, we performed phylogenetic and evolutionary studies to analyze the codons that might be under positive selection pressure and to estimate the time of the most recent common ancestor (T_MRCA_) of Brazilian YFV 2017 samples.

## RESULTS

### YFV samples geographic distribution and genome characterization

From February to June 2017, we collected 15 YFV samples from five distinct infected host species including mosquitoes, NHP and humans from 11 localities belonging to three main river basins across the current epidemic/epizootic territory in the Atlantic coast in ES and RJ (Table 1; Fig. 1). We determined and analyzed the whole YFV genome of twelve samples: two pools of infected mosquitoes belonging to two species *Haemagogus leucocelaenus* and *Hg. janthinomys*, six from NHP (four howler monkeys and two marmosets), and four from human cases (Table 1). Partial sequences of the YFV genome were also obtained from an additional human case (H189) (data not shown).

**Table 1.**
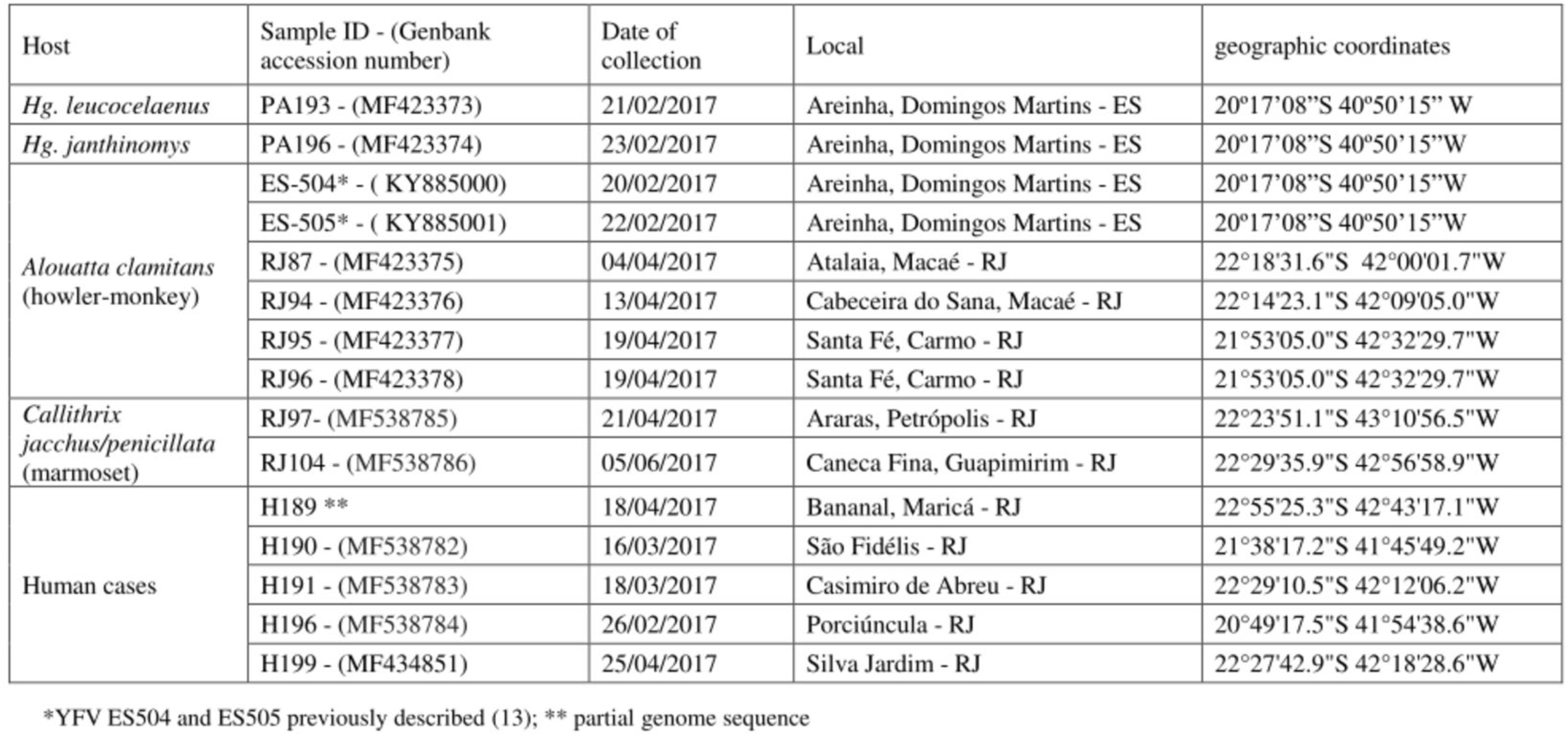
YFV samples collected in the 2017 Brazilian outbreak

**Figure 1.**
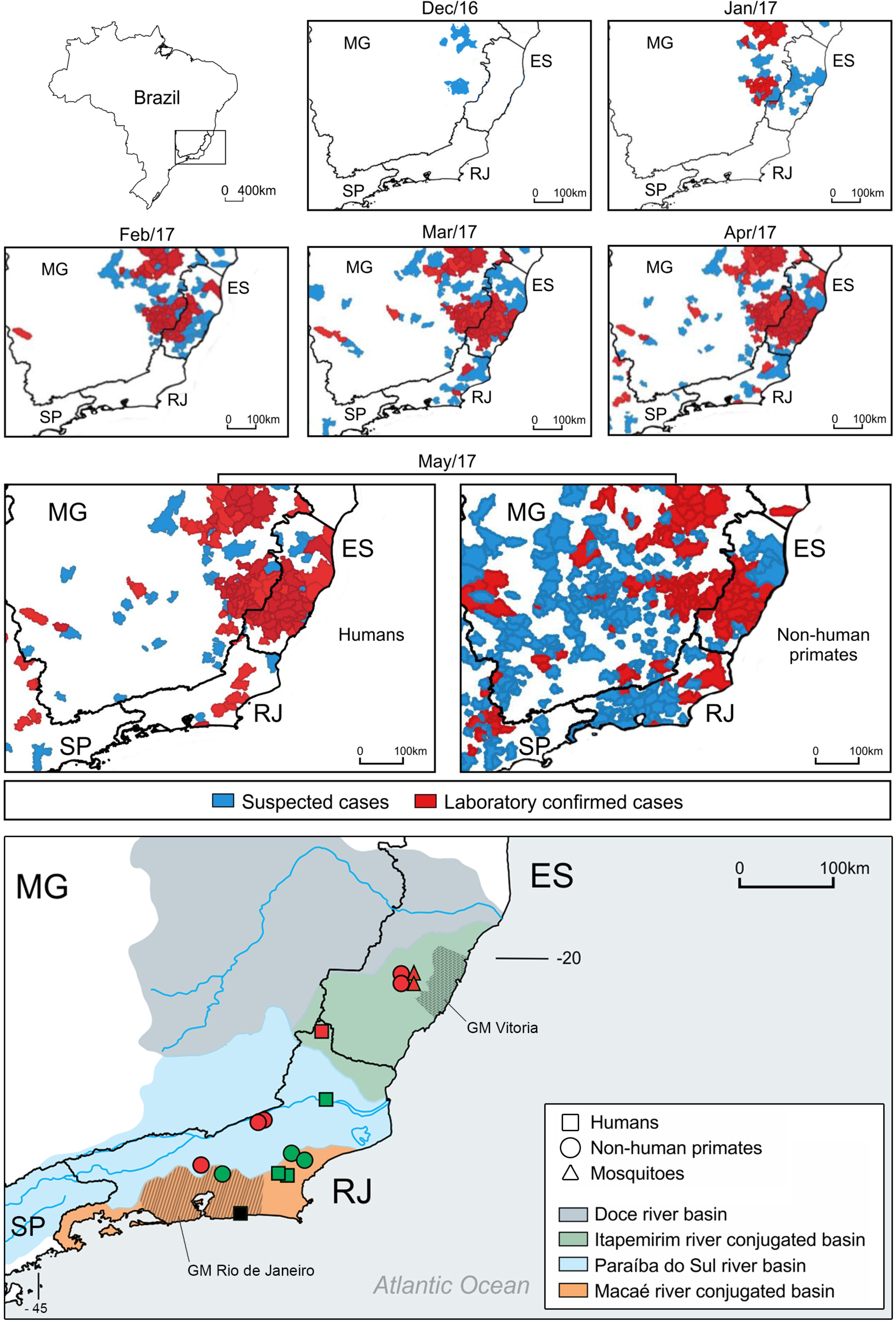
The spatiotemporal spread of the yellow fever outbreak in Southeast Brazil from December 2016 to May 2017, and geographical origins of the yellow fever virus (YFV) samples according to states, river basins, hosts, and YFV sub-clades A (red) and B (green). The black square corresponds to the H189 sample whose only partial sequences of the YFV genome were obtained. Brazilian states: ES (Espírito Santo), MG (Minas Gerais), RJ (Rio de Janeiro) and SP: (São Paulo). Hatched areas correspond to the Great Metropolitan (GM) areas of Rio de Janeiro and Vitória.

The comparison of all the YFV genomes from the ongoing Southeast Brazilian outbreak reveals low genetic variation, providing a mean nucleotide identity of 99.8 % and amino acid identity values ranged from 99.9% to 100%. However, part of the nucleotide variations is non-synonymous, leading to new amino acid substitutions in the polyprotein sequence (Fig. 2; Supplementary file 1). Regardless the host, all the 2017 YFV Brazilian genomes display a set of eight unique amino acid substitutions, that we have recently identified in two YFV samples from infected howler monkeys (ES-504 and ES-505) from ES state (15). Remarkably, the comparison with the genome of other South America strains confirmed that these polymorphisms are only present in the Brazilian strains from the ongoing outbreak when comparing with other South American strains. They localize at C protein (V108I), at NS3pro (E1572D; R1605K), at NS5 in MTase domain (K2607R; V2644I; G2679S), and at NS5 in RdRp domain (V3149A; N3215S) (Fig. 2). Nevertheless, the partial nucleotide sequence from H189 also displayed all amino acid changes detected in the other sequences, except for the mutation in the C protein. Moreover, all 2017 YFV Brazilian strains also share an amino acid change at position N/D2803S that was previously observed only in a Venezuelan strain isolated in 2006 (GenBank, KM388818). We also detected additional amino acid substitutions, which are not present in all the 2017 Brazilian genomes (Fig. 2). Accordingly, the YFV H199 sequence shows a change from an alanine (A) to a serine (S) at the amino acid position 826, corresponding to the NS1 protein (position 48). The YFV genomes RJ95, RJ96, RJ97, H191 and PA193, show a substitution from an isoleucine (I) to a valine (V) at position 2176 (NS4A, position 77). The phylogenetic analysis of prM/E sequences indicates that all 2017 YFV Brazilian strains cluster inside sub-lineage 1E of the Modern lineage of South America genotype I in a monophyletic clade with high support (bootstrap = 87 %) (Fig. 2 – Fig. Supplement 1). The same clustering pattern was obtained when we performed the phylogenetic analyses of either NS3 or NS5 nucleotide sequences (Fig. 2 – Fig. Supplement 2).

**Figure 2.**
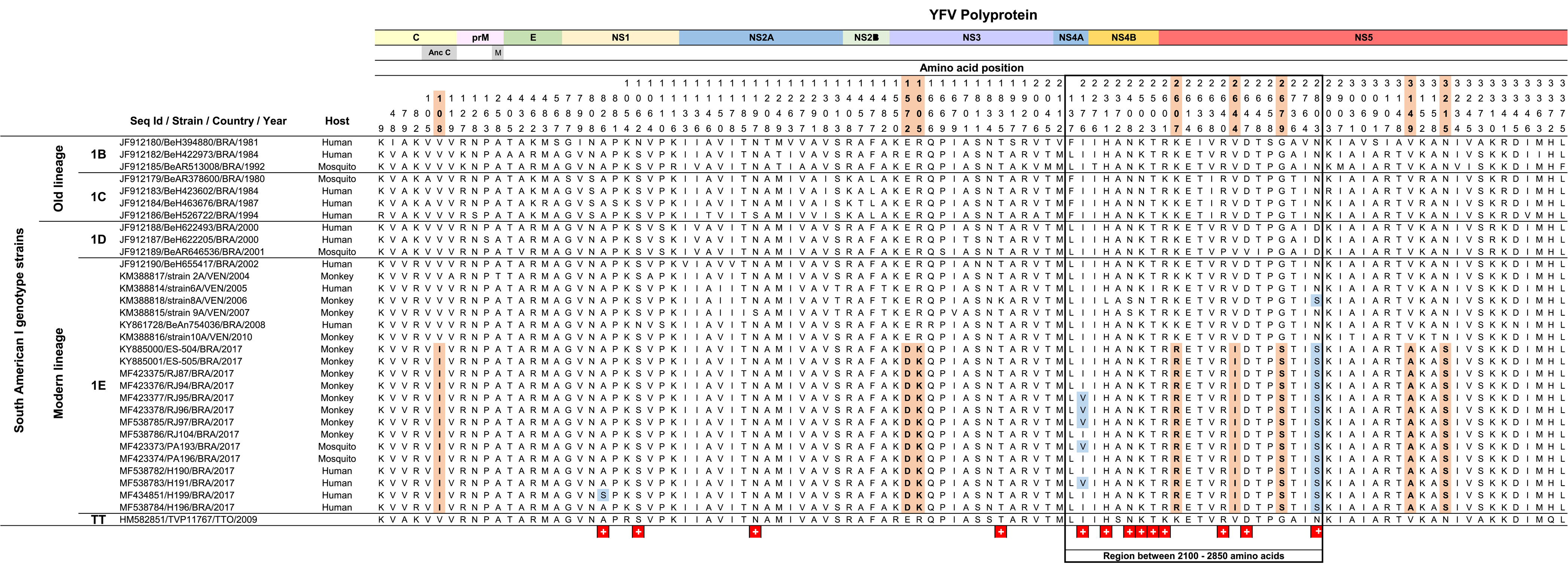
Amino acid (aa) differences revealed by the alignment of the precursor polyproteins of 32 yellow fever (YF) viruses of the South America genotype I. On the left of the alignment data, the identification of lineage, sub-lineages and yellow fever virus (YFV) sequences are supplied. On the top of the alignment, the YF viral proteins positions are indicated along with the aa positions of amino acid differences. The orange-highlighted aa indicates the position of aa shared only by all YF sequences from the ongoing outbreak in Brazil. Amino acid residues highlighted in blue indicate aa changes present in YF strains from the current outbreak, first described in the current study. The “+” symbol indicates the sites under positive selection in the YF polyprotein.

### Modeling and Structural Analysis of NS2B/NS3 and NS5 proteins

We created structural models of NS3 and NS5 proteins to gain some insights into the structural and functional effects of these amino acid substitutions. Initially, prior to the structural model generation, we aligned the NS3 and NS5 amino acid sequences of the prototype 2017 Brazilian YF virus (strain ES505) (15) with the different template sequences (Fig. 3– Fig. Supplements 1; Fig. 4– Fig. Supplements 1).

The effect of the amino acid substitutions on both NS3 and NS5 proteins was evaluated through hydrogen bond formation and electrostatic analysis. The two substitutions found in NS3, E88D and R121K (polyprotein position: E1572D; R1605K, respectively) are conservative and, as such, they have little impact on the surface electrostatic potential (Fig. 3 C, D). For the E88D substitution, a small structural change was observed, which mainly consisted of lysine 174 (polyprotein position: K1658) side chain displacement due to the loss of a hydrogen bond with the protein backbone (Fig. 3 E, F). On the other hand, the R121K substitution is located near the NS2B binding groove and might influence the interaction between these two molecules. Hydrogen bond analysis indicates that K 121 could favor the formation of a hydrogen bond with threonine 77 of NS2B (polyprotein position: T1431), whereas such an interaction was not identified in 2010 model (Fig. 3 A, B). This interaction could, in turn, modulate the NS3-NS2B binding affinity and, thus, the protease efficiency.

**Figure 3.**
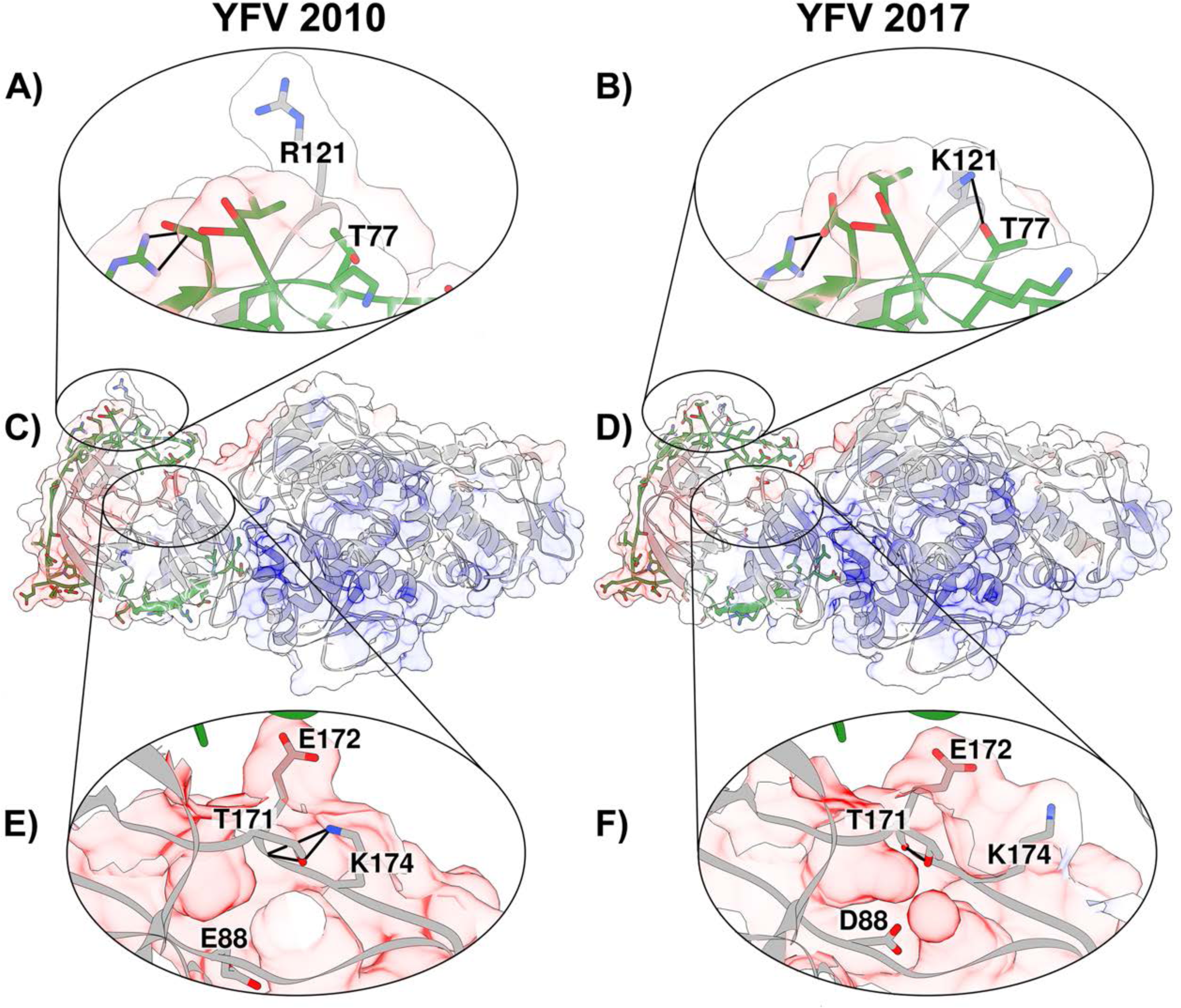
Tridimensional models obtained by comparative modeling of the NS2B-NS3 protein complex. Cartoon and surface electrostatic potential representation of R121K substitution (A, B), whole complex (C, D), and E88D substitution (E, F). The molecular surface is colored according to electrostatic potential, where red, white and blue correspond to acidic, neutral and basic potentials, respectively. NS2B is shown in green. Thick black lines represent hydrogen bonds (A, B, E, F)

The three first amino acid substitutions in NS5 are clustered in the MTase domain, whereas the remaining ones are in the RdRp domain. All amino acid substitutions at the MTase domain are conservative, but they are spatially adjacent. Arginine 101 (polyprotein position: R2607) alpha carbon is 9.7 Å away from that of isoleucine 138 (polyprotein position: I2644), which in turn is 10.6 Å away from serine 173 alpha carbon (polyprotein position: S2679). These three residues face the RdRp domain tunnel opening (Fig. 4B), which presents a basic electrostatic profile to accommodate the YFV RNA molecule (Fig. 4 – Fig. Supplement 2), suggesting that they may influence the enzyme activity.

**Figure 4.**
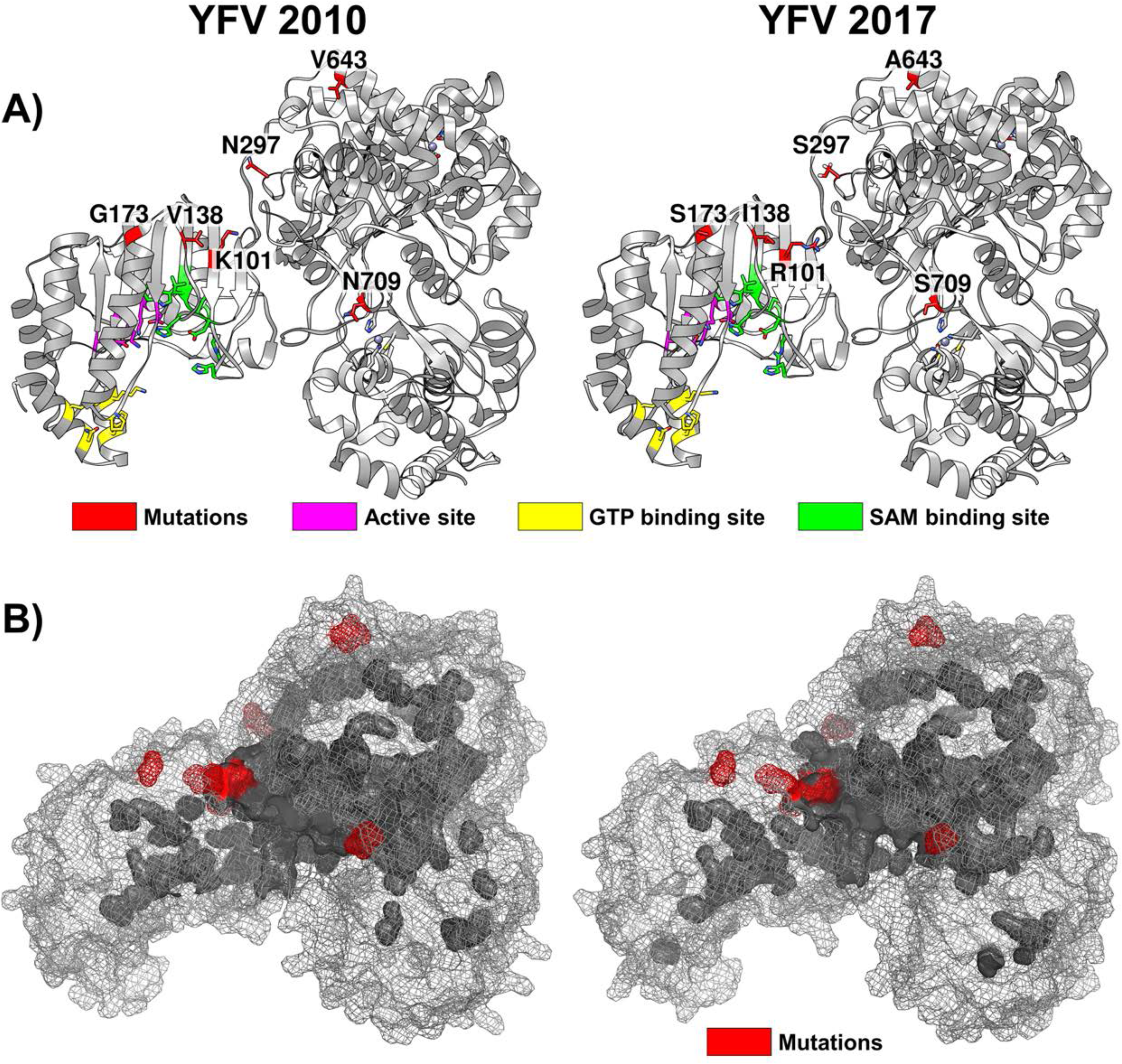
Tridimensional structural models obtained by comparative modeling of NS5 protein. (A) Cartoon representation of NS5 protein. Amino acid substitutions and binding site residues are shown in the sticks and colored according to the legend. (B) Cavities of NS5 protein. Amino acid substitution sites are shown in red.

Additionally, the N297S substitution (polyprotein position: N/D/S 2803) from the RdRp domain, although being conservative, is located near the hinge domain. The remaining two amino acid alterations: V643A (polyprotein position: V/A 3149) and N709S (polyprotein position: N/S 3215) are located at the protein surface and are also conservative. The combined effect of the mutations on the NS5 protein dynamics was assessed through molecular dynamics simulations, which indicate that the NS5 protein from 2017 sample has a decrease in fluctuation around the hinge region (Fig. 4– Fig. Supplement 3).

### Selection analyses

The mean dN/dS ratio of substitutions per site estimated by the SLAC method for the South America genotype I (SAI) and West Africa (WA) data sets was 0.05 and 0.04, respectively; thus indicating that purifying selection was the main evolutionary force of both YFV genotypes. Tests for negative/positive selection, however, reveal some important differences in the evolutionary dynamics of both genotypes (Fig. 5).

**Figure 5.**
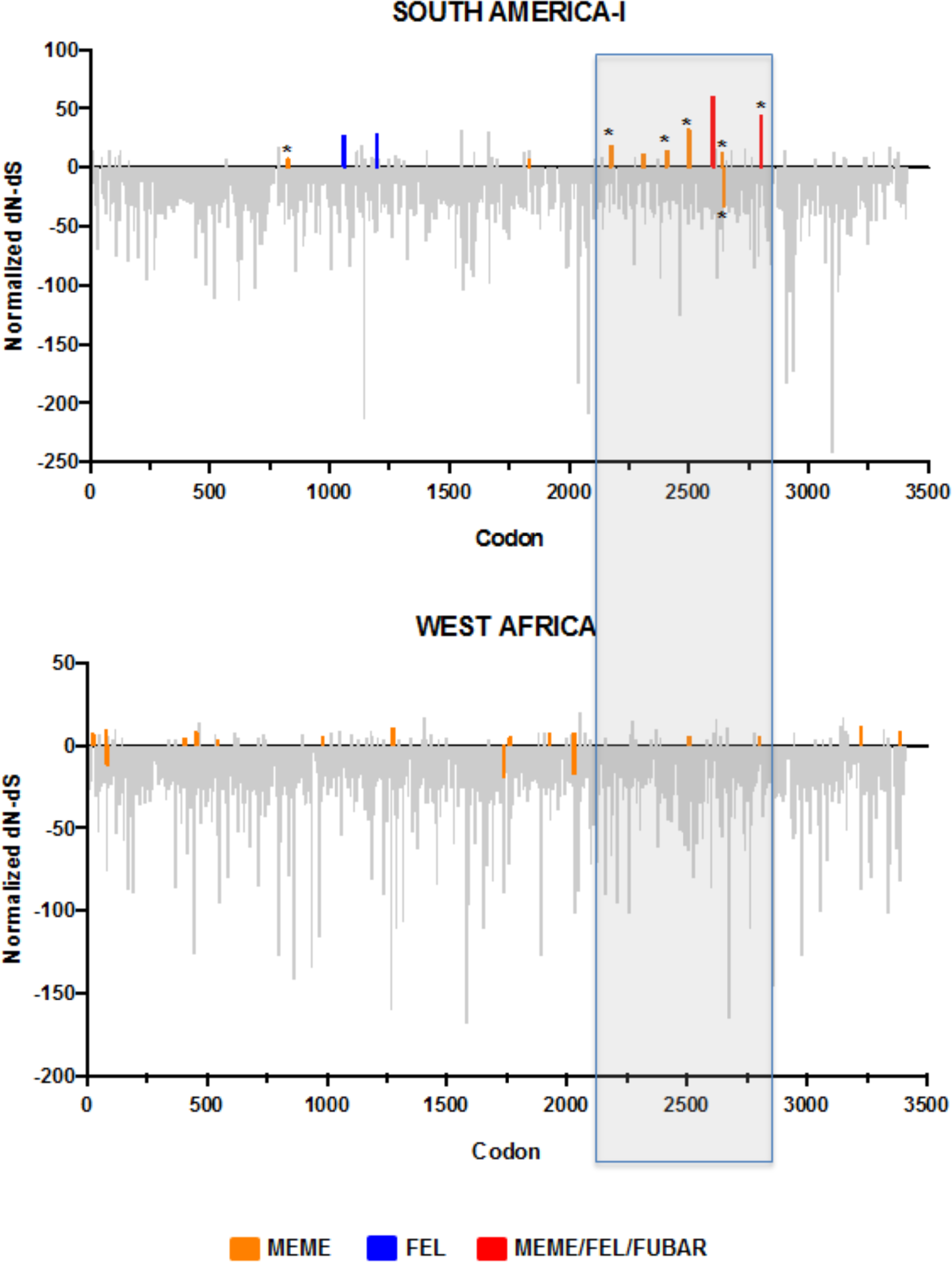
Codons of the yellow fever virus under positive selection in South America genotype I (top), and West Africa genotype (bottom). The Y-axis represents normalized dN-dS (non-synonymous substitutions minus synonymous substitution), and the X-axis represents codon positions. The region between codon positions 2100 and 2850 is highlighted inside a gray rectangle. Positively selected sites are shown with the color code that appeared at the bottom of the figure.

Selection analysis of the SAI dataset identifies 13 codons with evidence of positive selection, including nine sites with evidence of episodic diversifying selection (MEME), four sites with evidence of pervasive diversifying selection (FEL and/or FUBAR) and one site identified by all three algorithms (FEL, MEME, and FUBAR). Most sites (69%, 9/13) under positive selection were concentrated in a short genome segment (2,100-2,850 codon positions) coding for non-structural proteins NS4A (I2176V), NS4B (H2311L, N2408S, K2502N, T2503I) and NS5 (R2601K, R2640P, D2647V, N/D2803S) (Fig. 5 – Fig. Supplement 1). Selection analysis of the WA dataset, by contrast, detected no sites under pervasive diversifying selection and identified 18 sites with evidence of episodic diversifying selection (MEME) uniformly distributed along structural and non-structural proteins (Fig. 5 – Fig. Supplement 1).

### Evolutionary analyses

The Bayesian MCC tree inferred from the complete coding sequence (CDS) of YFV South American genotypes I and II reveals that YFV strains from the ongoing Southeast Brazilian outbreak grouped in a highly supported (*Posterior Probability* [*PP*] = 1) monophyletic cluster nested within sub-lineage E strains of the modern-lineage (Fig. 6). We further observed that 2017 YFV Brazilian strains segregate in two reciprocally monophyletic subgroups, one sub-cluster comprising strains of mosquitoes, NHP and humans from RJ and ES states (sub-clade A) (*PP* = 0.29), and one highly supported (*PP* = 0.99) sub-cluster containing YFV strains of NHP and humans from the state of RJ (sub-clade B). The median substitution rate of YFV South American genotypes complete genomes was estimated at 3.5 x 10^-4^ subst./site/year (95% HPD: 2.4-4.8 x 10^-4^ subst./site/year), in agreement with previous estimations(12). The median T_MRCA_ for all Brazilian 2017 YFV strains was estimated in April 2016 (95% HPD: July 2015 - October 2016) and for the sub-clade B at October 2016 (95% HPD: April 2016 - January 2017). In addition, the median T_MRCA_ of Venezuelan and Brazilian 2017 YFV strains was estimated as occurred 24 years ago.

**Figure 6.**
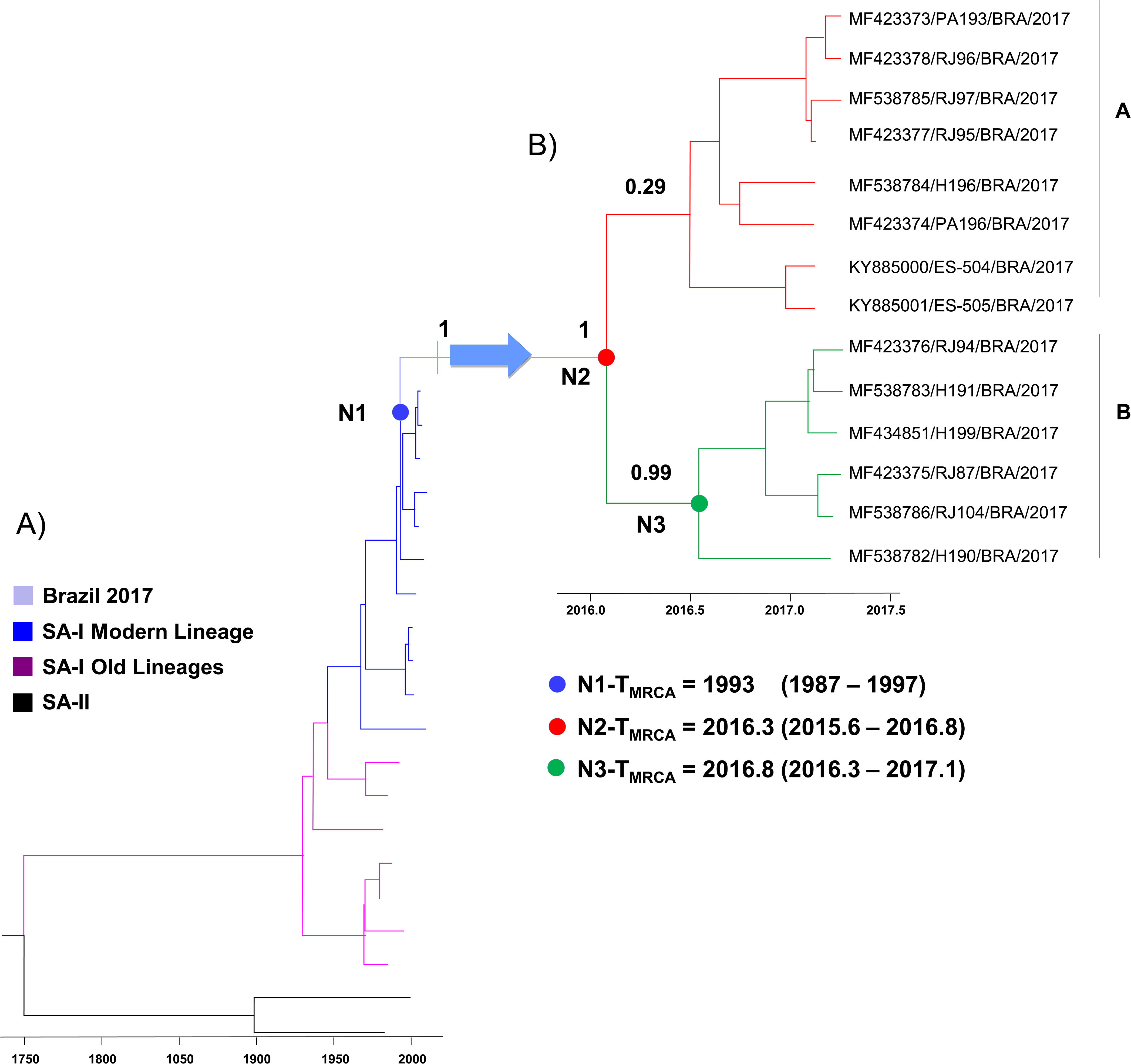
Phylogenetic evolutionary analysis based on the yellow fever virus (YFV) complete coding region. (A) Time-scaled Bayesian MCC tree of YFV CDS. The color code is explained at the left of the figure. Names and accession numbers of the strains are shown in Fig. 6 – Fig. Supplement 1. (B) YFV from the ongoing Southeast Brazilian outbreak.

## DISCUSSION

In the 2016-2017 YFV outbreak in Southeast Brazil, most of the epizootics and human cases originally occurred in inland rural areas of MG and subsequently in Eastern Atlantic coastal sites under the influence of the ES rain forest segments. After that, the YFV spreading approached and even touched the Great Metropolitan areas of Vitoria (ES) and Rio de Janeiro (RJ), where lived nearly 1.8 and 12.3 million unvaccinated inhabitants, respectively. Alarmingly, these densely populated areas host some of the busiest South American airports, ports and road networks, are highly infested by YFV competent urban vectors, *Aedes aegypti* and *Ae.albopictus,* and have repeatedly been the territory of severe dengue epidemics. All together, these ecological and sanitary marks raised concern about the potential risk of YFV to reemerge in an urban cycle in Brazil as well as to spread to other countries and continents rapidly(23, 24). Furthermore, YFV strains characterized by a set of amino acid polymorphism were identified during this outbreak, and their biological impact needed to be investigated (15).

Regardless of deriving from five distinct host species infected in a six month lag in 11 sites dispersed along 350 Km of the Southeast Brazilian coast across the outbreak territory including the Great Metropolitan area of Rio de Janeiro, the YFV samples analyzed in the current study are almost identical at the nucleotide and amino acid levels. Also, they share a molecular signature represented by nine amino acids, being eight in highly conserved positions at NS3 and NS5 encoding regions, and one in the structural capsid protein. Previous analysis of 2017 YFV from two howler monkeys from a single site in ES had not considered the substitution at position 2803 at NS5. However, in the current study, the analysis of a higher number of samples allowed the identification of one more substitution at position 2803 at NS5 in all 2017 YFV. This molecular signature represented by nine amino acids have never been observed before in South-American and African genotypes (15).

Hypothetically, amino acid changes at conserved protein positions of NS3 and NS5 may have a role in the capacity of viral infection to vertebrate and invertebrate hosts and thus accelerating the spreading of the ongoing outbreak. The NS3 and NS5 proteins have multiple enzymatic activities essential for viral RNA replication and 5´-capping (25-27). For these reasons, we also investigated the impact of the identified amino acid substitutions through the structural analysis of NS3 and NS5 protein models. Even though all the amino acid substitutions were mainly conservative, they occur close to domains that might be affected by these subtle modifications. Furthermore, we were able to detect features that may correlate with an increase in enzymatic efficiency and constitute an advantage in viral dissemination. In the NS3 protein, the R121K substitution is located in the region responsible for the interaction with NS2B, the cofactor for the proteolytic activity of this enzyme. Although both residues bear a positive charge, lysine was shown to potentially establish a hydrogen bond with NS2B, due to the less bulky side chain. For the NS5 protein, we found that the three amino acid substitutions located at the MTase domain were spatially contiguous and could influence the relative orientation between the two domains. The N297S substitution might also have a significant role in the enzymatic efficiency since is located near the hinge domain. Hence, molecular dynamics simulations have shown that all these substitutions combined may have a stabilizing effect on the linker domain, which has been demonstrated to influence the enzyme processivity directly and viral replication in Dengue 4 virus *in vitro* models (28, 29). These findings shall be addressed in future studies considering the unrevealed diversity of the YFV in the cell culture and animal models.

A hot spot region of episodic or pervasive positive selection was identified in between codons 2100-2850 of CDS region in YFV genomes of South American genotype I, that was not detected in the West Africa genotype. Interestingly, most of these positively selected sites localize in the NS4B, and NS5 coding regions have also been described in Zika virus from the current epidemics(30). Several studies have demonstrated the role of non-structural proteins in the host innate immune response against flavivirus infection (22, 31-33). Onward, these proteins widely interact with other viral proteins and host molecules (27, 33-35). It is also interesting to note that the 2803 position, part of the molecular signature of 2017 YFV, is one of the positively selected sites. Also, two other positions positively selected (826 and 2176), presented variability in the 2017 YFV. These observations support some singularities in the evolutionary dynamics of YFV South American genotype I and also indicate that fixation of some amino acid substitutions in Brazilian 2017 YFV strains might have been driven by positive selection.

Since the beginning of the XXI, a striking spreading of the YFV has been occurring in Brazil. The outbreaks formerly constrained to the endemic Amazon and Central-Western regions have reached the South and Southeast of Brazil, where vaccination coverage against YF was insignificant until the explosion of the ongoing outbreak. It has been recently proposed that a YFV strain from Trinidad-Tobago introduced in Brazil and Venezuela in the late 1980s originate all modern-lineages strains belonging to sub-lineages 1D and 1E (12). The main source of variability related to the ancestral lineages is the occurrence of several amino acid substitutions particularly within non-structural viral proteins (12), as observed in the 2017 Brazilian outbreak YFV. All Brazilian 2017 YFV belonged to the sub-lineage 1E and clustered with the Venezuelan strains isolated in the late 2000s, consistent with the notion that ancestral YFV strain responsible for the ongoing Brazilian outbreak would have originated in Venezuela (12).

Although Brazilian 2017 YFV strains are most closely related with Venezuelan 2004-2010 YFV strains, they share a relatively distant common ancestor traced to 1993 (95% HPD: 1987-1997) (Fig. 6). This indicates that the virus may have circulated in endemic South American regions for a long period before being introduced in Southeast Brazil, but the precise pathway of viral dissemination is difficult to elucidate because the paucity of Brazilian YFV sequences sampled from endemic regions over the last 10-15 years. We estimate the median T_MRCA_ for the Brazilian 2017 YFV strains at early 2016, suggesting that the virus circulated for several months in the Southeastern region before the ongoing outbreak was first recognized at December 2016. This pre-detection period of cryptic transmission of YFV in Southeastern Brazil is comparable to that recently estimated for Zika virus in the Northeastern Brazilian region (36).

Bayesian analysis also showed that Brazilian 2017 YFV strains segregate into two sub-clusters, one of them (sub-clade A) containing sequences from both RJ and ES, and the other (sub-clade B) comprising only sequences from RJ whose T_MRCA_ was traced to October 2016. It is also interesting to note that all YFV sequences of RJ that branched within sub-clade A were sampled from sites situated in the Paraíba do Sul basin whose tributaries born on the northern side of the Serra do Mar, a 1,500km long system of mountain ranges and escarpments that runs parallel to the Atlantic Ocean coast in Southeastern Brazil (Fig. 1). By contrast, YFV sequences of RJ that branched within sub-clade B were obtained from locations situated along the Macaé conjugated river basin whose tributaries born on the escarpments of the coastal side of the Serra do Mar.The only exception is the sample H190, which despite originating in the Paraíba do Sul river basin, it clustered in the sub-clade B. Intriguingly, the sampling location of H190 (São Fidelis) is located in the largest discontinuity of the long mountain ranges of the Serra do Mar and this YFV strain branched basally to all other sub-clade B strains. Together, these results support the occurrence of multiple independent introductions of YFV in RJ (probably from both ES and MG) and further indicate at least two main routes of viral dissemination within the state running at the northern and coastal sides of the Serra do Mar. These analyses also point that YFV dissemination through the coastal route in Rio de Janeiro probably started in São Fidelis around late 2016, but more YFV sequences from other Southeastern states are necessary to confirm this conclusion.

Future analysis based on reverse genetic approaches can contribute to establishing the role of amino acid substitutions present in Brazilian 2017 YFV in the viral fitness and transmissibility. It will also be crucial to improving the number of YFV genomes from Brazilian endemic and non-endemic regions in the last years to better understand the epidemiology during recent years.

## MATERIAL AND METHODS

### Ethics Statements

This study was reviewed and approved by the Ethics Committee for human research at the Instituto Oswaldo Cruz (IOC) (CAAE 69206217.8.0000.5248), which exempted the need of a specific written informed consent from patients or their legal representatives. The protocols for mosquito rearing as well as handling and blood sampling of NHP were approved by the Institutional Ethics Committee of Animal use at IOC (CEUA licenses LW-34/2014 and L037/2016, respectively). Capture of wild NHPs and mosquitoes were approved by the Brazilian environmental authorities: SISBIO-MMA licenses 54707-137362-2 and 52472-1, and INEA license 012/2016012/2016. No specific permits are needed for conducting mosquito collection in the urban and suburban areas in Southeastern Brazil. This study does not include endangered or protected species.

### Mosquito samples

Adult mosquitoes were collected with BG-sentinel adult traps baited with dry ice as a source of CO_2_ as well as with an insect net when approaching to humans in the forest, at the forest fringe, and in the modified environment. The insects were immediately frozen in dry ice or N2, transported to the laboratory, identified to species according to Consoli and Lourenço-de-Oliveira (36), and pooled according to species, the number of captured mosquitoes and collecting site and day. Entire bodies of pooled mosquitoes were ground and treated in Leibovitz L15 medium (Invitrogen) supplemented with 4% Fetal Bovine Serum (FBS), and submitted to the RNA extraction as described elsewhere(37).

### Non-human primate samples

Blood samples were taken from the femoral vein of dying NHPs or the cardiac cavity of recently dead NHPs. Samples from howler monkeys were centrifuged (2,000 *g* for 10 min) for obtaining plasma or serum samples stored at low temperature (N_2_ or – 80 °C) until RNA extraction. Due to the small amount of blood in the cardiac cavity of dead marmosets, RNA extraction was performed from total blood frozen in dry ice immediately after collection in the field.

### Human samples

Blood samples were obtained from patients for diagnosis procedures made at their respective municipal public health ambulatories. Serum samples of suspicious clinical YF cases were sent to the State Central Laboratory in Rio de Janeiro (LACEN-RJ) for molecular and serological analyses. Aliquots of serum samples of YFV laboratory-confirmed cases and negative for Zika virus, Chikungunya virus and Dengue virus infection were selected by the LACEN-RJ staff and stored at - 80 °C until RNA extraction.

### YFV RNA extraction, screening for YFV infection by RT-PCR and Nucleotide Sequencing

YFV RNA from mosquito homogenates, blood, and serum samples were obtained by using the QIAamp Viral RNA kit (Qiagen). The RNA samples were eluted in 60 μL of AVE buffer and stored at - 80 ° C until use. The YFV RNA was reverse transcribed using the Superscript IV First-Strand Synthesis System (Invitrogen) with random hexamers or specifics primers at 25 °C for 10 min, 55 °C for 10 min and 80 °C for 10 min. Detection of YFV genome was performed by conventional PCR as described elsewhere (15). Complete YFV genome amplification was carried out by conventional PCR using KAPA HiFi PCR kit (KAPABIOSYSTEMS) under the following conditions: 95 °C for 3 min, followed by 35 cycles at 98°C for 20 sec, 65°C for 15 sec and 72°C for 45 sec, and a final extension at 72 °C for 1 min 30 sec. The set of primers utilized in PCR and sequencing procedures are listed in the Supplementary files 2 and 3, respectively. Complete genome sequences were deposited in the GenBank database (Table 1). Amplified products were sequenced as previously described (15). The sequenced amplicons were analyzed, and contigs were assembled by using SeqMan Pro v8.1.5 (3), 414 (DNASTAR, Madison, WI). The sequences were manually edited and compared with other sequences available in GenBank (https://www.ncbi.nlm.nih.gov/genbank/). The Molecular Evolutionary Genetics Analysis (MEGA) 7.0 program was used to calculate nucleotide and amino acid distances, as well as to explore the amino acid signatures observed in the YFV strains from the ongoing outbreak in Southeast Brazil.

### Comparative modeling, optimization and MD simulation of NS3 pro and NS5 proteins

We performed the modeling of the NS2B-NS3 protein complex and NS5 protein of the 2017 outbreak YFV prototype (Genbank, KY885001) and the 2010 Venezuelan 10A strain (Genbank, KM388816). Initially, we used BLASTP program (https://blast.ncbi.nlm.nih.gov/Blast.cgi?PAGE=Proteins), defining the Protein Data Bank (PDB) as a search set, to select the template structures for comparative modeling. Three PDB structures were used as templates for the NS2B-NS3 protein complex. Template 2VBC (51% identity and 96% of coverage) corresponds to the crystal structure of the NS3 protease-helicase from dengue virus(38). Template 1YKS (96% identity and 70% of coverage) comprises the NS3 helicase domain of the YFV(39). Template 5GJ4 comprises the NS3 protease domain (56% identity and 27 % of coverage) and NS2B peptide cofactor (42% identity and 31 % of coverage) of Zika virus(40). For the NS5 model generation, a single PDB structure from Japanese Encephalitis virus (4K6M) (19) was used as a template, which shares 60% identity with the YFV sequence. Template and target sequences were then aligned using the PSI-Coffee mode of T-Coffee program(41). One hundred homology models were generated using the standard “auto model” routine of Modeller version 9.18 (42) for each target sequence. Each model was optimized using the variable target function method (VTFM) optimization until accomplishing 300 iterations. Molecular dynamics (MD) optimization, in the slow level mode, was carried out. The full cycle was repeated two times to produce an optimized conformation of the model. The resulting modeled structures were selected according to their discrete optimized protein energy (DOPE) score. The GROMACS version 5.1.2 package(43), was used to carry out minimization using the AMBER99SB ILDN force field(44). A short minimization procedure of 150 steps (100 steps of steepest descent + 50 steps of conjugate-gradient) was performed. Initial and optimized models were evaluated by DOPE, Ramachandran plot and QMean server (Supplementary File 4) (45). The electrostatic potential analysis was conducted using the APBS program(46). Atomic partial charges and atomic radii parameters from the Amber force field were assigned using the PDB2PQR server (47). Figures of sequence alignments were rendered using ALINE (48), and 3D structures were generated using UCSF Chimera and PyMol.

Molecular dynamics simulations were carried out using the GROMACS package, with the AMBER99SB-ILDN force field. Protonation states were assigned using pdb2pqr software, and the zinc-coordinating cysteine residues were manually deprotonated. The models were further optimized prior to the MD runs, through 5.0 x 10^6^ steps of the Steepest Descent (with and without heavy atom restraints) and Conjugate Gradients algorithms. The systems were then run, under an NPT ensemble, for 500 ps with restraints and 2.0 x 10^5^ ps without restraints. The V-rescale thermostat and Berendsen barostat were used for temperature and pressure control, respectively. The 2010 and 2017 strains and replicas were simulated at 297 and 310 K for a total of 8 runs and 1.6 x 10^6^ ps. Analysis was made over the final 150 ns of the production runs.

### PrM/E phylogenetic analysis

A 666-nucleotide sequence consisting of the last 108 nucleotides of the prM gene, the entire M gene (225 nucleotides), and the ?rst 333 nucleotides of the E gene was used to established genotype the YFV strains, as previously described(5). In addition to the sequences obtained in the current study, a set of sequences of the prM/E junction fragment were selected using the Blast tool (https://blast.ncbi.nlm.nih.gov/Blast.cgi) and all sequences were aligned using Molecular Evolutionary Genetics Analysis (MEGA) 7.0(49). The phylogenetic tree was generated by the Neighbor-joining method (50) using a matrix of genetic distances established under the Kimura-two parameter model (51). The robustness of each node was assessed by bootstrap resampling (2,000 replicates) (52). The homologous region (prM/E) of a dengue virus strain available at the GenBank database (AF349753) was used as an outgroup.

### Selection pressure analyses

Two datasets of sequences were determined: a) Dataset South America I (Set SAI) with all complete CDS of South American YFV stains available in the GenBank plus sequences obtained in the current study (N=32); b) Dataset West Africa (Set WA), with complete CDS from West Africa genotype YFV strains available in the Genbank. The reason to generate these two datasets was that West African genotype has a closest genetic relationship with South America genotype I (5), and only two CDS from YFV strains belonging to South America genotype II are available in the GenBank to date. Datasets were aligned using Bioedit v7.2.3 (53) and were tested for positive selection by using the online adaptive evolution server DATAMONKEY(54, 55). The ratio of non-synonymous to synonymous substitutions (dN/dS) per codon sites were estimated using four different methods, SLAC - Single Likelihood Ancestor Counting, FEL - Fixed Effects Likelihood(56), MEME - Mixed Effects Model of Evolution(57) and FUBAR - Fast Unbiased Bayesian Approximate(58). The analysis was run based on neighbor-joining tree and significant P-value (< 0.1). The automated model selection tool available in the server was used for selection of appropriate nucleotide substitution bias model for both datasets.

### Evolutionary analyses

Complete coding regions sequences (CDS - 10,239 nt in length) of YFV of American origin (South America genotypes I and II), with a known date of isolation, were obtained from the GenBank database (http://www.ncbi.nlm.nih.gov). Retrieved sequences were aligned together with sequences obtained in the current study using MEGA 7.0 (49). The nucleotide substitution rate and time of the most recent common ancestor (T_MRCA_) of American YFV strains were estimated using the Markov chain Monte Carlo (MCMC) algorithm implemented in the BEAST v1.8.3 package(59, 60) with BEAGLE (61) to improve run-time. The best-fit nucleotide substitution model, a relaxed uncorrelated lognormal molecular clock model(62), and a Bayesian Skyline coalescent tree prior(63) were used. The uncertainty of parameter estimates was assessed after excluding the initial 10% of the run by calculating the Effective Sample Size (ESS) and the 95% Highest Probability Density (HPD) values, respectively, using TRACER v1.6 program(64). TreeAnnotator v1.7.5 (60) and FigTree v1.4.0 (65) were used to summarize the posterior tree distribution and to visualize the annotated Maximum Clade Credibility (MCC) tree, respectively.

## Acknowledgments

To Monique Albuquerque Motta for the help on mosquito identification; Iule de Souza Bonelli, Marcelo Quinteka Gomes, Lidiane S. R. Menezes, Nathália D. Furtado, Stephanie O. D. da Cruz and Adalgiza as Silva Rocha for technical assistance; Heloisa Diniz and Leônidas Leite Santos for the help with illustrations; Alexandre Otávio Chieppe, Ana Paula Martins Brandão, Carlos Augusto Ferandes (Secretaria Estadual de Saúde do Rio de Janeiro) Alessandro Pecego M. Romano (Grupo Técnico de Vigilância de Arboviroses) and Roberta Gomes de Carvalho (Programa Nacional de Controle da Dengue) Brazilian Ministry of Health, Gilsa Aparecida P. Rodrigues (Secretaria de Saúde do Estado do Espírito Santo), Gilton Luiz Almada (Centro de Informação Estratégica de Vigilância em Saúde-ES) for the access to epidemiological data and support for the field work; Marilza L. Lange (Secretaria Municipal de Saúde, Municipality of Domingos Martins), Luciano L. Salles (Vigilância em Saúde, Municipality of Ibatiba) Agenor Barbosa de Oliveira, Marcelo dos Santos Rosário, Mauro Cesar Louzada, Max Mauro dos Santos Coelho for their assistance in field collections in Espírito Santo. Alexandre Bezerra de Souza and Vicente Klonowski (Atalaia Park), Romenique de Lemos Araújo, Luiz Ribeiro Nogueira and Raquel Giri Faria (Environmental Guard) and Macaé prefecture for their support in collecting samples. Raul Henrique Rafael and Ana Luiza Quijada for access to the Guapimirim sample; Cassia Schitino de Carvalho Gomes and Paulo Vitor Araújo Gomes for access to the Carmo samples.

### Competing interests

The authors declare that no competing interests exist.

## Figures –Figure Supplement legends

**Figure 2 - Figure Supplement 1.**
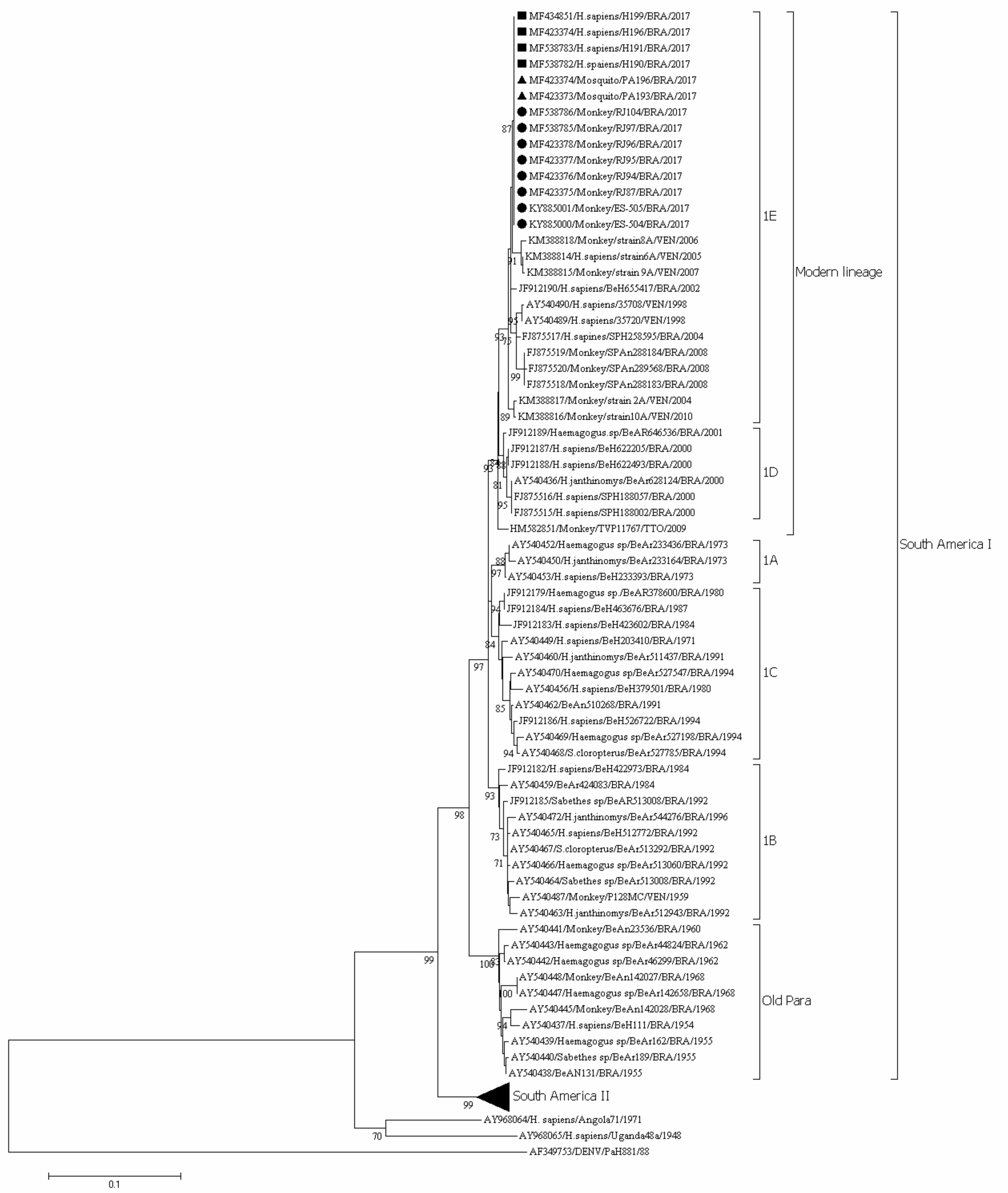
Phylogenetic analysis based on the prM/E junction region of yellow fever virus (YFV) strains analyzed in the current study and YFV sequences retrieved from the National Centre for Biotechnology Information (NCBI). Only bootstrap values up to 70% are shown. YFV genotypes are shown at the right side of the figure. The scale bar at the bottom represents 0.1 substitutions per nucleotide position (nt.subst/site). YFV from the 2017 ongoing Southeast Brazilian outbreak are marked with a filled triangle (mosquito strains), filled square (human strains) and filled circle (non-human primates strains).

**Figure 2 - Figure Supplement 2.**
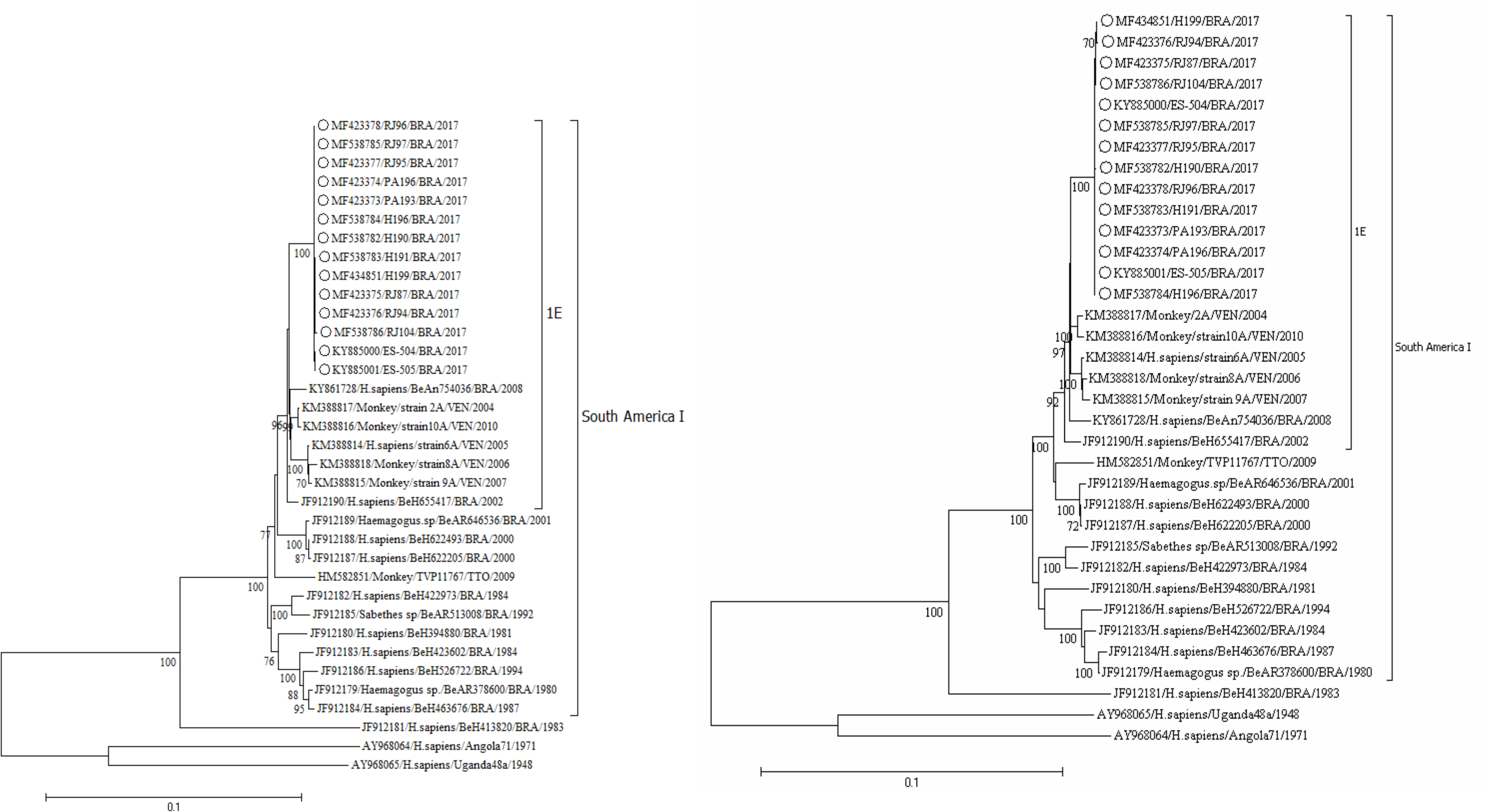
Phylogenetic analysis based on the NS3 (A) and NS5 (B) encoding region of yellow fever virus (YFV) strains analyzed in the current study and 21 YFV sequences retrieved from the National Centre for Biotechnology Information (NCBI). Only bootstrap values up to 70% are shown. The scale bar at the bottom represents 0.1 substitutions per nucleotide position (nt.subst/site). YFV from the 2017 ongoing Southeast Brazilian outbreak are marked with an empty circle. South America genotype I and sub-clade 1E are shown at the right side.

**Figure 3 - Figure Supplement 1.**
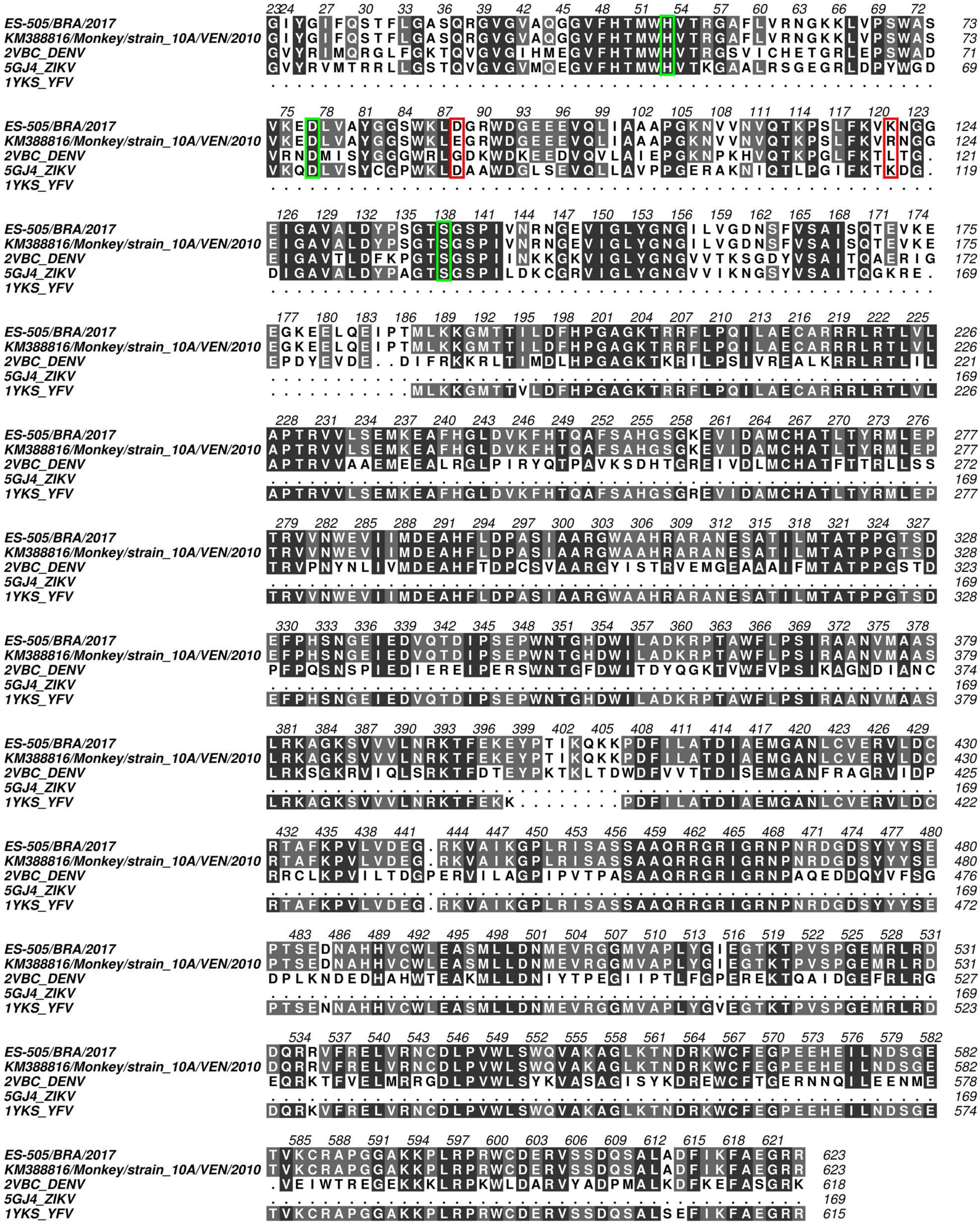
Multiple sequence alignment of the NS3 protein sequence belonging to the 2017 Brazilian yellow fever virus (strain ES505), 2010 Venezuelan 10A strain and the three templates used in comparative modeling experiment. Black and gray filled positions of the alignment represent fully and partially conserved residues, respectively. Red outline highlights the amino acid found in the 2017 Brazilian strain. Green outline highlights the positions of the active site residues.

**Figure 4 - Figure Supplement 1.**
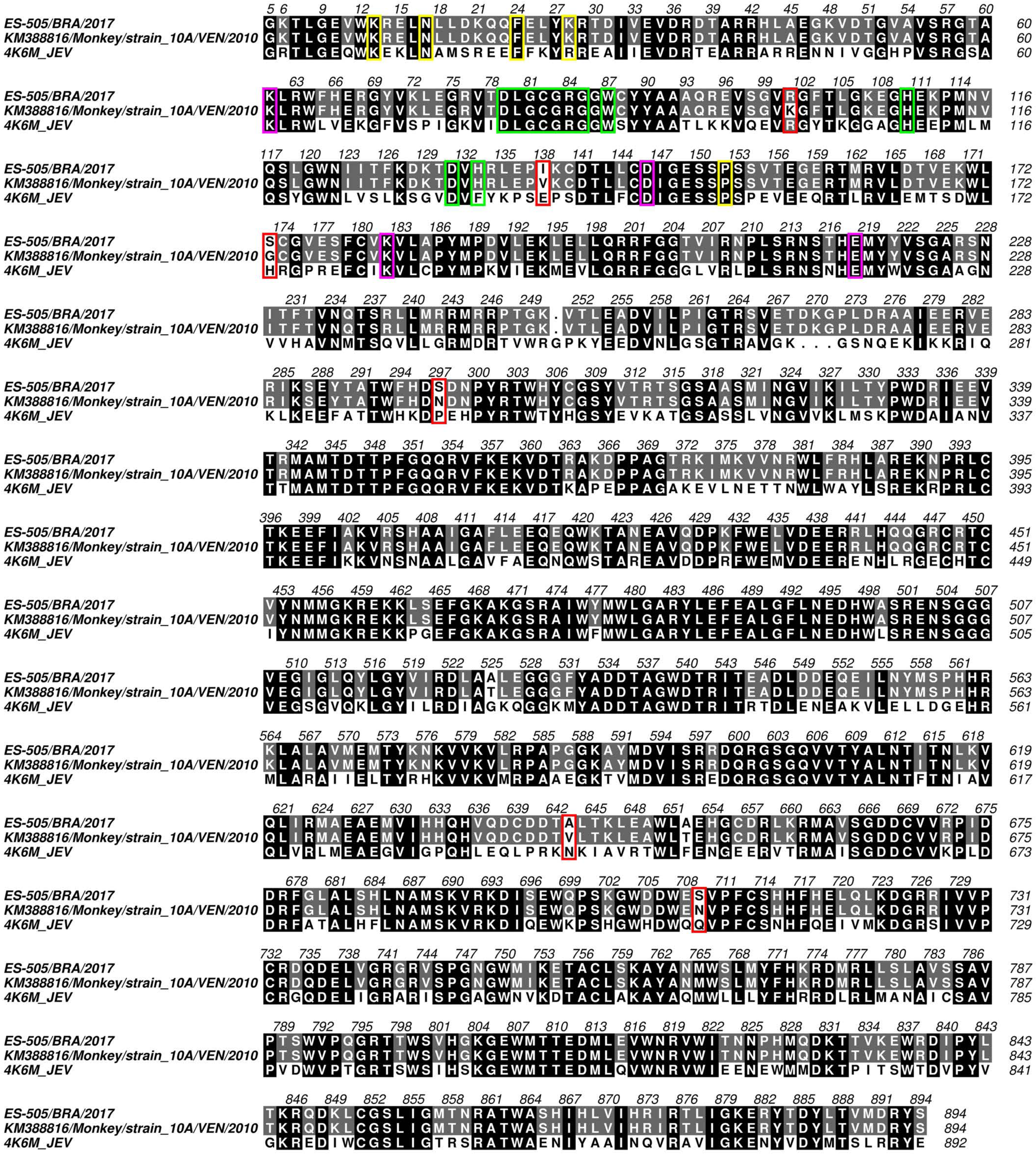
Multiple sequence alignment of the NS5 protein sequence belonging to the 2017 Brazilian yellow fever virus (strain ES505), 2010 Venezuelan 10A strain and the template used in comparative modeling experiment. Black and gray filled positions of the alignment represent fully and partially conserved residues, respectively. Red outline highlights the amino acid found in the 2017 Brazilian strain. Pink, yellow and green outlines highlight the positions of residues found in the active site, GTP binding site, and SAM binding site, respectively.

**Figure 4 - Figure Supplement 2.**
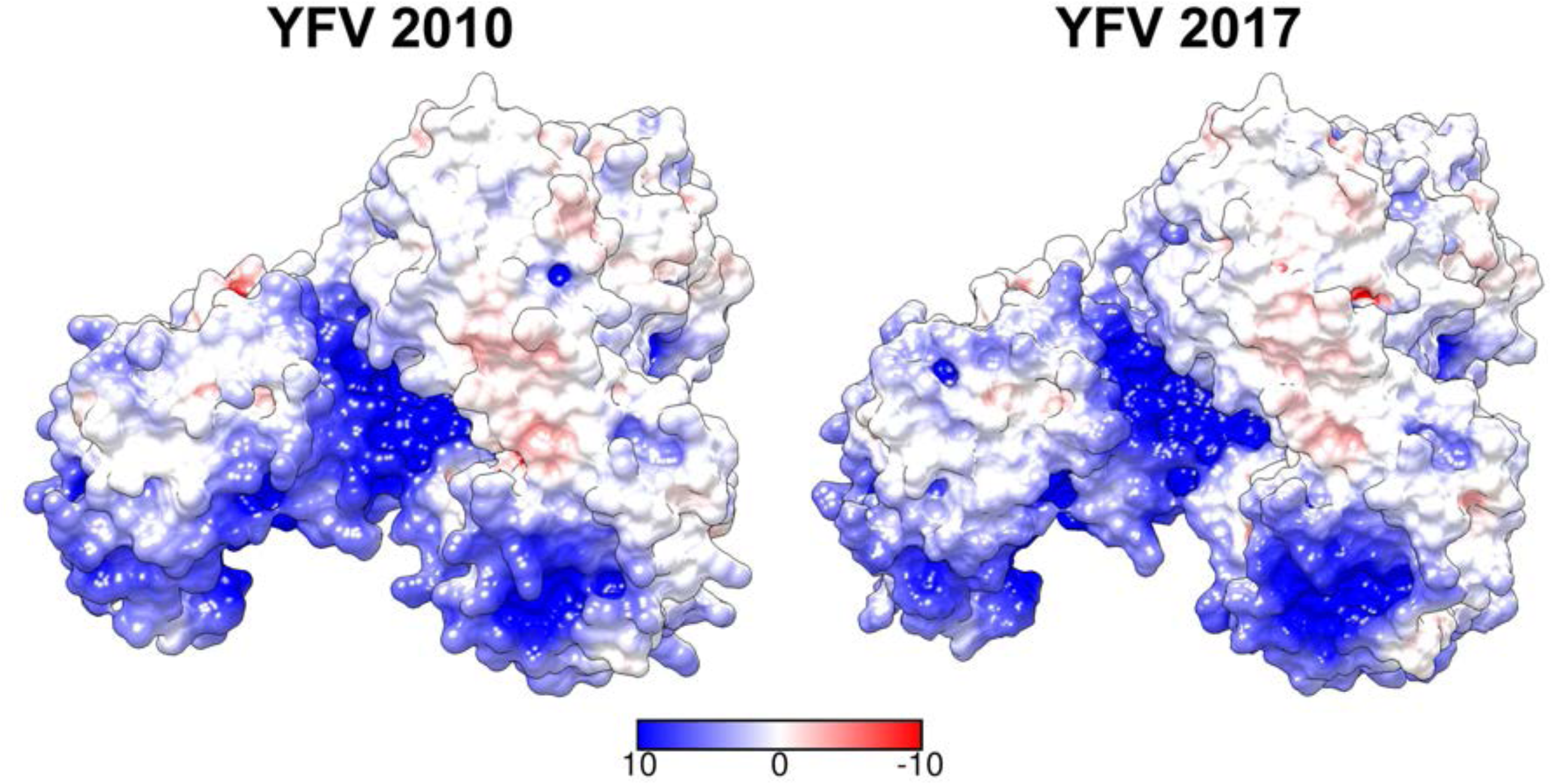
The surface electrostatic potential of NS5 protein. The molecular surface is colored according to electrostatic potential, where red, white and blue correspond to acidic, neutral and basic potentials, respectively.

**Figure 4 - Figure Supplement 3.**
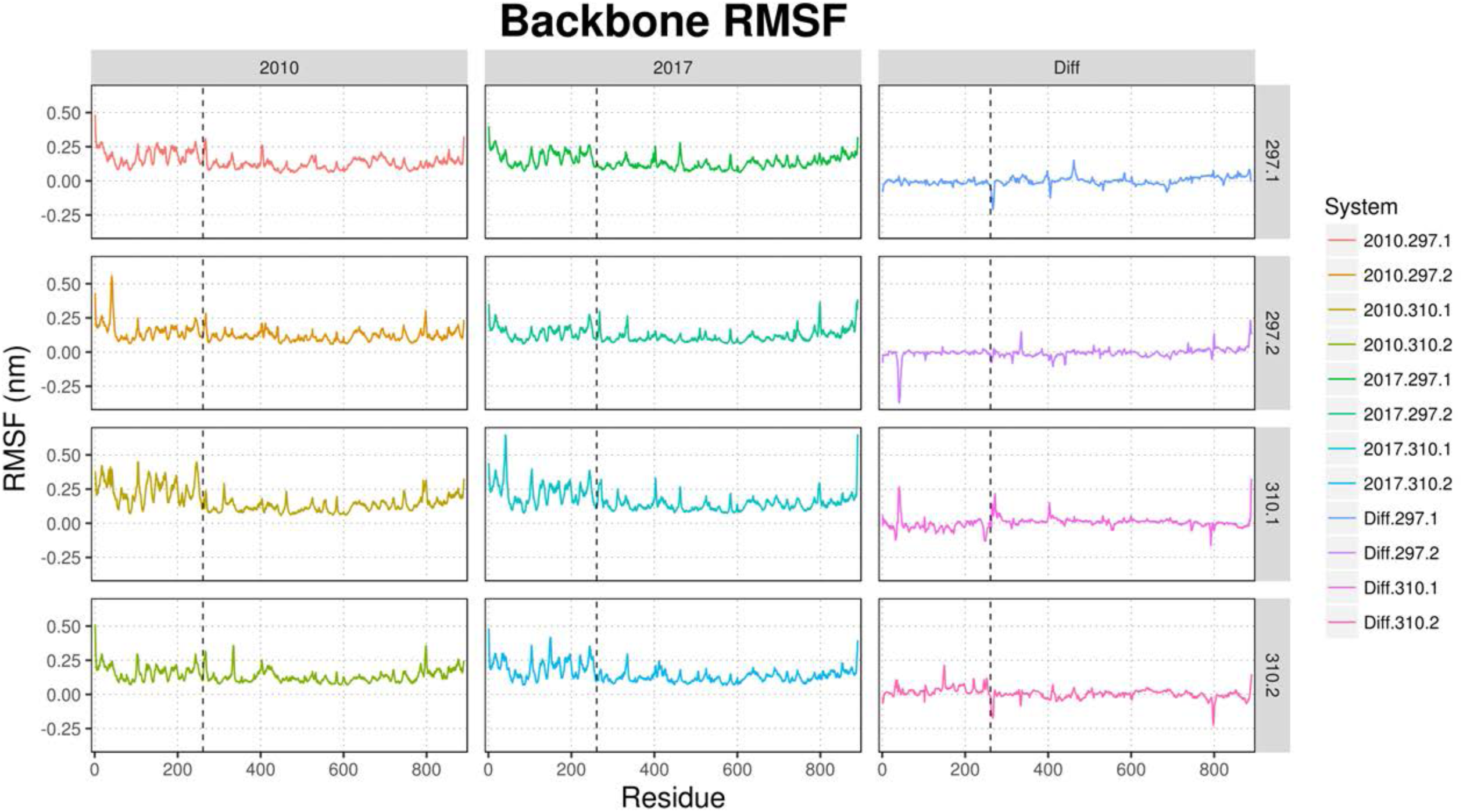
Root mean square fluctuation plots of the final 150 ns of each NS5 MD production run. The systems were run in replicates with distinct random seeds for initial velocities generation. Temperatures of 297 and 310 K were used. The last column (Relative) represents the subtraction of the calculated fluctuations of 2010 from the 2017 values. The vertical dashed line marks the hinge domain. Fluctuations were calculated with GROMACS software and plotted using R.

**Figure 5 - Figure Supplement 1.**
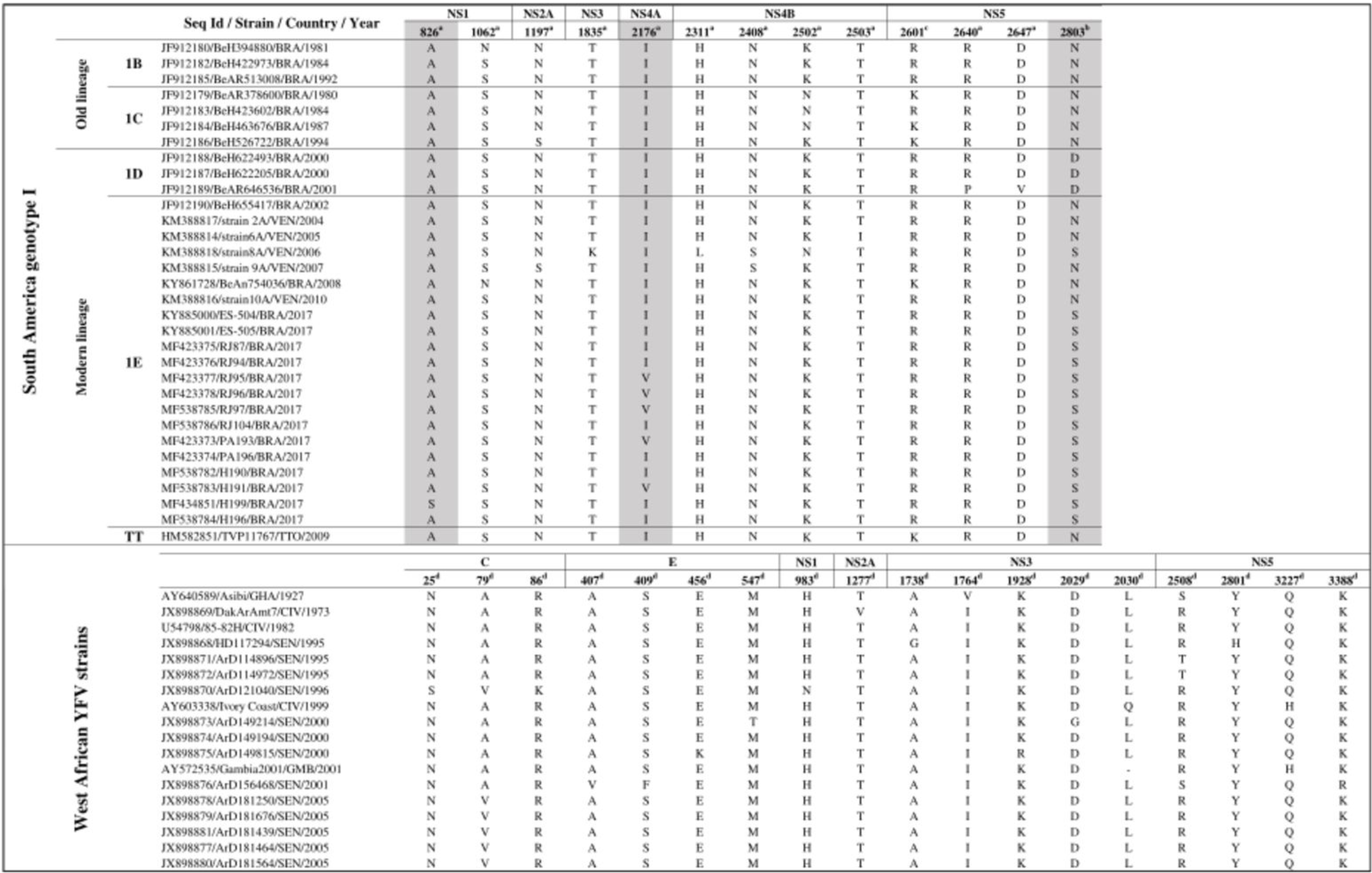
Positively selected sites in yellow fever virus (YFV) polyprotein from South American genotype I (top) and West African (bottom). Sites, where the non-synonymous substitution occurred in YFV 2017 (top) are shown with a gray shadow. a-Sites supported by one method (FEL or MEME); b-sites supported by two methods (MEME/FUBAR or fel/fubar); c-site supported by three methods FEL/MEME/FUBAR.

**Figure 6 – Figure supplement 1.**
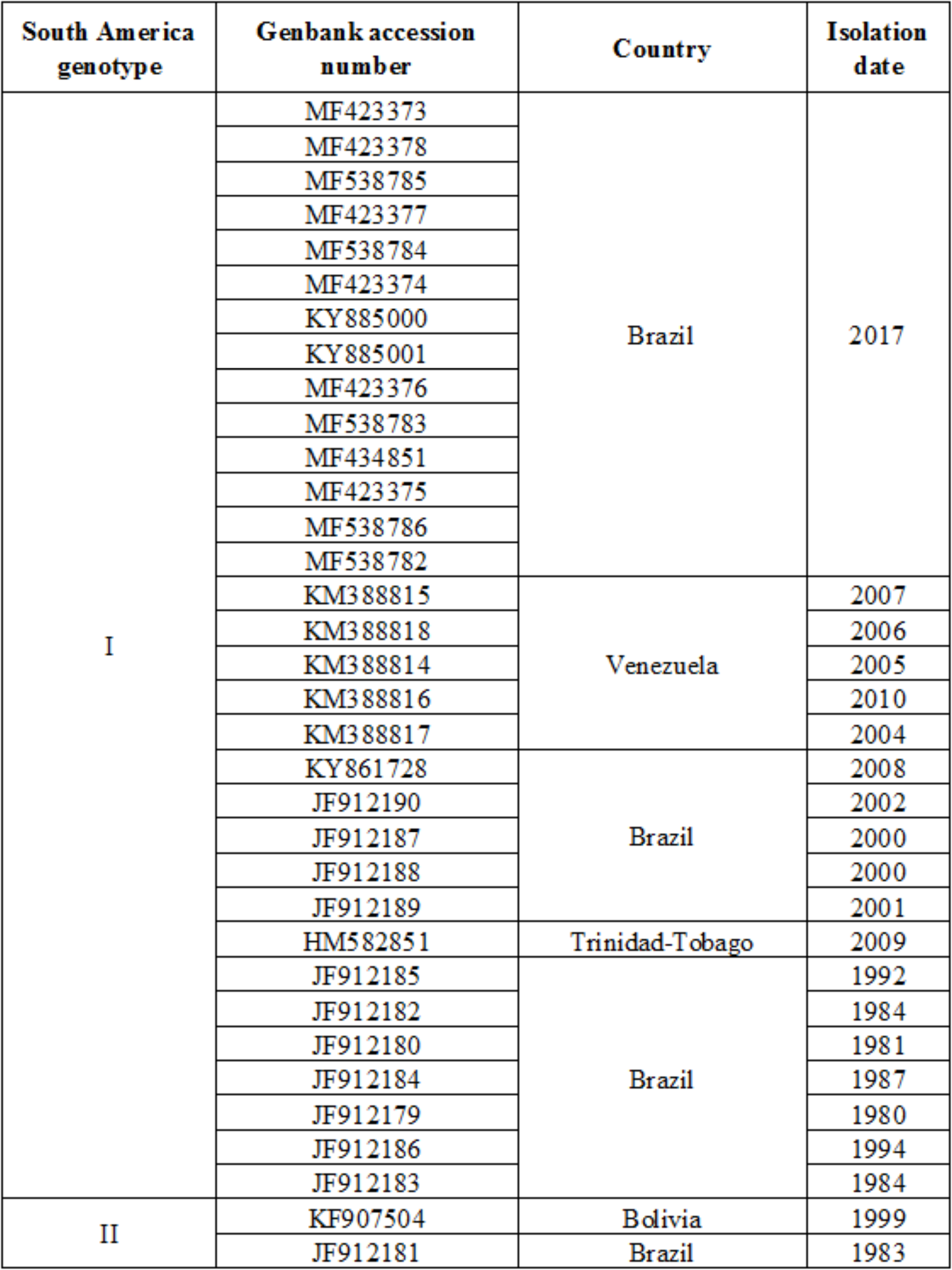
Yellow fever virus strains used in the Time-scaled Bayesian MCC tree.

